# Sex differences in lifespan trajectories and variability of human sulcal and gyral morphology

**DOI:** 10.1101/2020.10.02.323592

**Authors:** Covadonga M. Díaz-Caneja, Clara Alloza, Pedro M. Gordaliza, Alberto Fernández Pena, Lucía de Hoyos, Javier Santonja, Elizabeth E.L. Buimer, Neeltje E.M. van Haren, Wiepke Cahn, Celso Arango, René S. Kahn, Hilleke E. Hulshoff Pol, Hugo G. Schnack, Joost Janssen

**Affiliations:** Department of Child and Adolescent Psychiatry, Institute of Psychiatry and Mental Health, Hospital General Universitario Gregorio Marañón, Madrid, Spain; Ciber del Área de Salud Mental (CIBERSAM); Instituto de Investigación Sanitaria Gregorio Marañón (IiSGM), Madrid, Spain; School of Medicine, Universidad Complutense, Madrid, Spain; Departamento de Bioingeniería e Ingeniería Aeroespacial, Universidad Carlos III de Madrid, Madrid, Spain; Department of Psychiatry, UMCU Brain Center, University Medical Center Utrecht, Utrecht, The Netherlands; Department of Child and Adolescent Psychiatry/Psychology, Erasmus University Medical Centre, Sophia Children’s Hospital, Rotterdam, The Netherlands; Department of Psychiatry, Icahn School of Medicine at Mount Sinai, New York, United States

## Abstract

Sex differences in development and aging of human sulcal morphology have been understudied. We charted sex differences in trajectories and inter-individual variability of global sulcal depth, width, and length, pial surface area, exposed (hull) gyral surface area, unexposed sulcal surface area, cortical thickness, and cortex volume across the lifespan in a longitudinal sample (700 scans, 194 participants two scans, 104 three scans, age range: 16-70 years) of neurotypical males and females. After adjusting for brain volume, females had thicker cortex and steeper thickness decline until age 40 years; trajectories converged thereafter. Across sexes, sulcal shortening was faster before age 40, while sulcal shallowing and widening were faster thereafter. While hull area remained stable, sulcal surface area declined and was more strongly associated with sulcal shortening than with sulcal shallowing and widening. Males showed greater variability for cortex volume and thickness and lower variability for sulcal width. Across sexes, variability decreased with age for all measures except for cortical volume and thickness. Our findings highlight the association between loss of sulcal area, notably through sulcal shortening, with cortex volume loss. Studying sex differences in lifespan trajectories may improve knowledge of individual differences in brain development and the pathophysiology of neuropsychiatric conditions.

## Introduction

Females and males show disparate prevalence rates, age at onset and clinical manifestations of many psychiatric disorders and neurodevelopmental conditions (1). The study of sex differences in development of brain structure may offer valuable insights into their biological underpinnings and pathophysiology. Sexual dimorphism in brain morphology and function across development may be the consequence of genetic, molecular, hormonal, and early environmental influences, including but not limited to sex-specific differences in gene expression and cellular mechanisms, epigenetic programming, prenatal and postnatal hormone levels, and immune function (2). In humans, sex and gender also affect later exposure to and interaction with social and physical environments, in the context of gendered socialization processes in most cultures (2).

Previous literature assessing sex differences in human brain structure has shown larger total brain volume in males overall, with less consistent findings for specific regions, especially when results are adjusted for sex differences in total brain volume (TBV) (3). The largest cross-sectional study to date reported larger cortex volume and pial surface area in males and thicker cortex in females (4). However, once TBV was accounted for, reported sex differences in pial surface area and in developmental trajectories were no longer significant (3,4).

Pial surface area and cortical thickness are differentially associated with cognitive ability, are reported to have distinct genetic determinants, and between the two, surface area displays a stronger association with cortex volume (5–8). Pial surface area can be divided into unexposed, sulcal, surface area and exposed, gyral, surface area, which is also called hull surface area. Sulcal surface area in turn can be fractionated into sulcal depth, length, and width (9). In a seminal longitudinal study, sexually dimorphic trajectories were reported for unadjusted pial surface area and hull surface area, which peaked later in males between 3-30 years of age (10). Older males have larger sulcal width relative to older females, while no sex differences are present at younger ages, which may point to sex differences in age-related changes in sulcal width (11,12). Although cortical laminar architecture is different between gyri and sulci, assessment of gyral- and sulcal-specific morphology has been largely overlooked in longitudinal lifespan studies of sex differences in neurotypical individuals (13–15). Sulcal area and length are reportedly more strongly positively associated with brain volume than sulcal depth in young adolescents across sexes (16). However, the relative contribution of changes in gyral and sulcal metrics to loss of cortex volume and surface area across the lifespan in both sexes is still unclear (16).

The majority of studies assessing sex differences in cortex morphology have focused solely on assessing mean effects, which may be limited approach to appropriately address the evident overlap between male and female distributions and the high inter-individual heterogeneity among individuals of both sexes (4). Greater variance (the spread of the distribution) for cortical surface area and cortical thickness has been reported in males as compared with females (17,18). However, it is not clear if sex differences in inter-individual variability exist for sulcal morphological metrics and whether variability for these metrics is constant across the lifespan.

Here, we present a large longitudinal lifespan study assessing eight aspects of cortex morphology. The aim of this study was to chart developmental trajectories of cortex volume, pial surface area, sulcal surface area, hull surface area, cortical thickness, and sulcal depth, width, and length in females and males across the lifespan. We also assessed whether age and sex modulate the association between brain morphology and cortex size. Finally, we assessed whether sex influences the inter-individual variability of these eight measurements with age.

## Methods

### Sample

We included healthy individuals with at least two T1-weighted magnetic resonance imaging (MRI) scan acquisitions from a large longitudinal sample taken from two studies conducted in patients with schizophrenia and healthy participants. Detailed information regarding clinical assessments of the Utrecht Schizophrenia project and the Genetic Risk and Outcome of Psychosis (GROUP) consortium, Utrecht, The Netherlands, are described in (19–21). Exclusion criteria for healthy participants were the following: i) intelligence quotient (IQ) below 80, ii) major medical or neurological illness (including migraine, epilepsy, hypertension, cerebrovascular disease, cardiac disease, diabetes, or endocrine disorders), iii) past head trauma, iv) alcohol or other substance dependence, and v) presence of a lifetime diagnosis of psychotic disorder or a positive family history of a psychotic disorder in first- or second-degree relatives, as established with the Family Interview for Genetic Studies (22). The IRB at the University Medical Center Utrecht reviewed the study protocols and provided ethical approval. All participants provided written informed consent.

The selection of participants from the initial dataset is described in SFigure 1A. Exclusion of participants due to image quality assessment procedures is described in more detail in the Supplemental text and SFigure 1B. The final sample consisted of 700 scans, including at least two scans for 131 females and 167 males. Detailed demographic, cognitive and morphometric information for the final sample can be found in Table1.

**Table 1.**
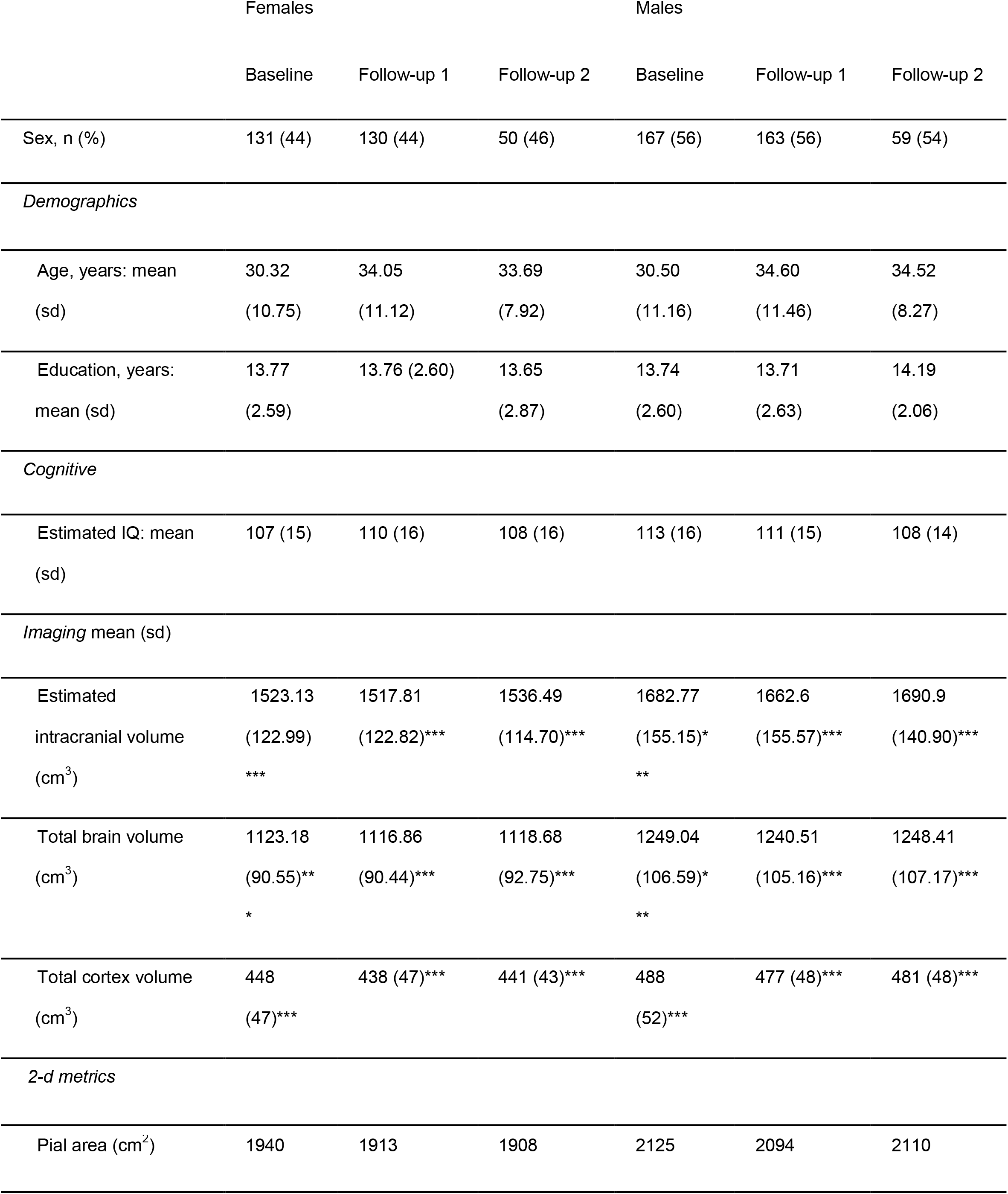
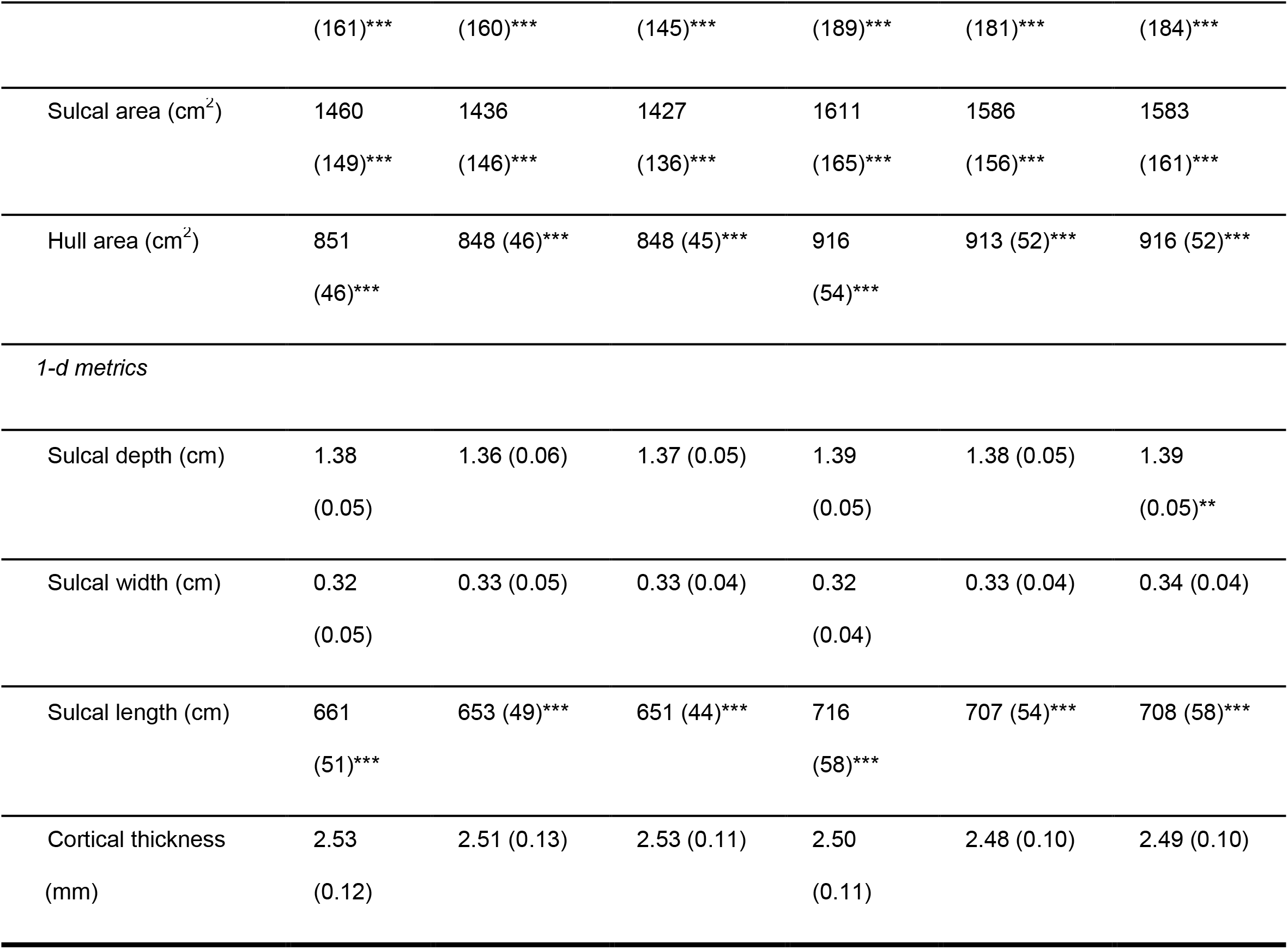
Demographic, cognitive and imaging characteristics of females and males. The total number of scans is 700. Significance calculated by Chi-square or Welch t-tests when appropriate. Significant sex differences at each of the timepoints are shown. *<0.05, **<0.01, ***<0.001. IQ, Intelligence quotient. Education and estimated IQ at baseline for participants with data available at each timepoint.

### acquisition and image analysis

Two scanners (same vendor, field strength and acquisition protocol) were used. Participants were scanned at least twice on either a Philips Intera or an Achieva 1.5 T and a T1-weighted, 3-dimensional, fast-field echo scan with 160-180 1.2 mm contiguous coronal slices (echo time [TE], 4.6 ms; repetition time [TR], 30 ms; flip angle, 30°; field of view [FOV], 256 mm; in-plane voxel size, 1×1 mm^2^) was acquired. All included participants had their baseline and follow-up scan/s acquired on the same scanner.

### Image processing

Global sulcal depth, width and length, pial surface area, exposed (hull) gyral surface area, unexposed (sulcal) surface area, and cortical thickness were assessed (see Figure 1A). All images were analyzed using the FreeSurfer analysis suite (v5.1) with default settings to provide detailed anatomical information customized for each individual (23). The FreeSurfer analysis stream includes intensity bias field removal, skull stripping, generation of a “ribbon” image and reconstruction of gray and white matter surfaces (24,25). Total brain tissue volume was derived as the sum of total gray and white matter volumes. Total cortical volume, total cortical surface area and mean global cortical thickness were then extracted from the FreeSurfer output. For all images, sulcal segmentation and identification was performed with BrainVISA software (v4.5) using the Morphologist Toolbox and Mindboggle software using default settings (9,26). After importing Freesurfer’s ribbon image into BrainVISA each sulcus is segmented with the cortical sulci corresponding to the crevasse bottoms of the “landscape,” the altitude of which is defined by image intensity. A spherical hull surface was generated from a smooth envelope that wrapped around the hemisphere but did not encroach into the sulci, a morphological isotropic closing of 6 mm was applied to ensure boundary smoothness. The median sulcal surface spans the entire space contained in a sulcus, from the fundus to its intersection with the hull. For each fold, sulcal area is defined as the total surface area of the medial sulcal surface, sulcal length is measured on the hull and is defined as the distance of the median sulcal surface intersecting the hull, and sulcal width is defined as the distance between each gyral bank averaged over all points along the entire median sulcal surface. FreeSurfer output was imported into Mindboggle software for calculating geodesic sulcal fundi depth. All metrics were measured in the native space of the participant’s images and left and right hemisphere values were either summed or averaged. FreeSurfer, BrainVISA and Mindboggle derived measurements have been validated via histological and manual measurements and have shown good test–retest reliability (27–29).

**Figure 1.**
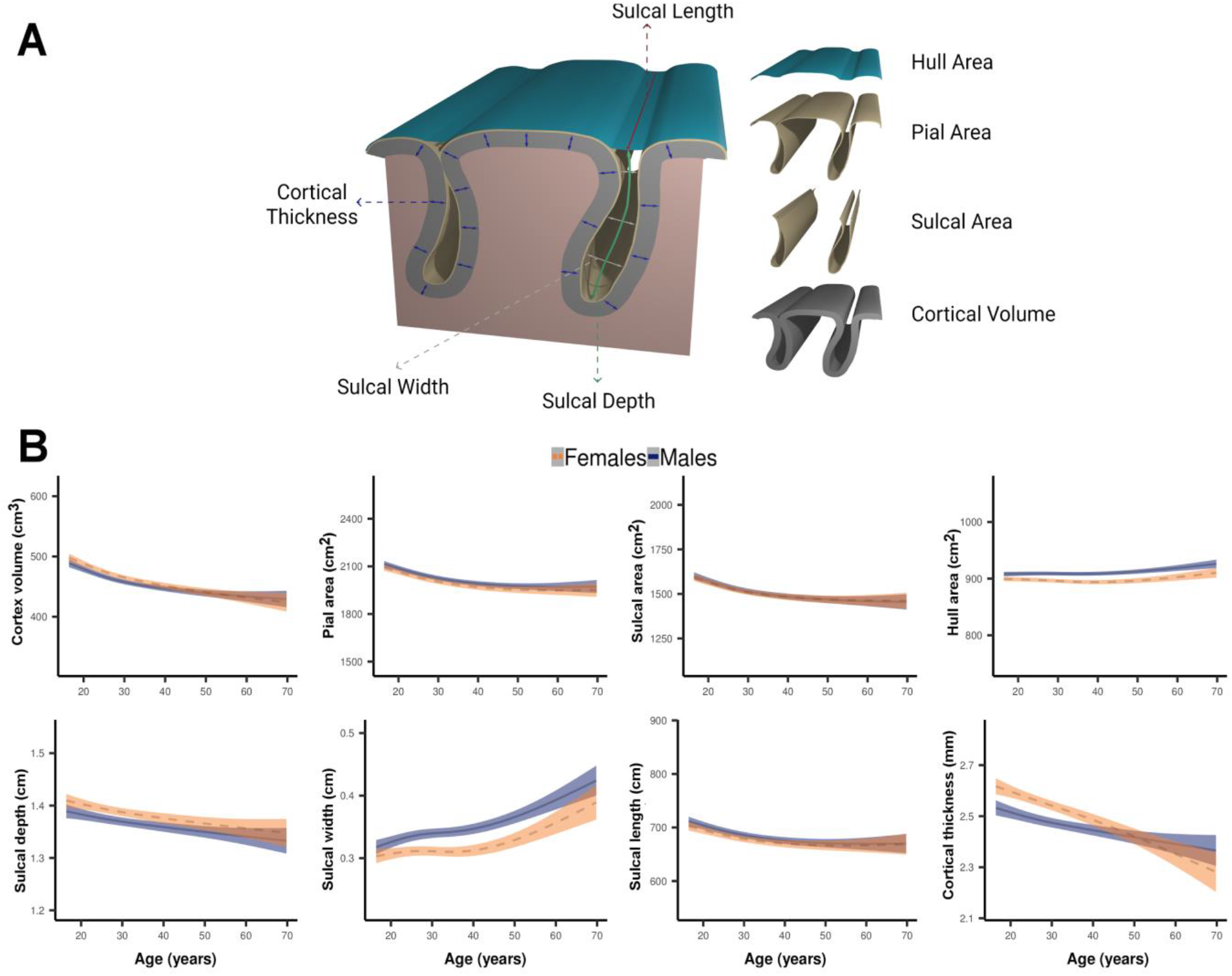
A: Schematic representation of the eight gyral and sulcal measurements. B: Age dependencies of the eight brain metrics for males (solid lines with 95% confidence intervals shown in blue) and females (dashed lines with confidence intervals in orange). Fits from generalized additive mixed models (GAMM). Total brain volume and scanner are included in the models as covariates.

### Statistical analysis

All analyses were performed in R (https://cran.rstudio.com/). In order to assess the amount of explained variance of the scanner on the morphological measurements, a visualization framework derived from gene expression analysis was adopted (http://bioconductor.org/packages/variancePartition) (30). As can be seen in SFigure 2, sex and age explained on average about 20% and 14% of the total variance, respectively. Scanner explained on average less than 1% of the variance at baseline and follow-up.

#### Longitudinal modeling of gyral and sulcal metrics: trajectories

To model the longitudinal trajectories of the brain metrics, generalized additive mixed models (GAMMs) were used (31). GAMMs are a generalization of Generalized Additive Models (GAMs) which are well suited for modeling nonlinear relationships through local smoothing on the response variable due to the interaction with independent covariates or complex relationships among them. In addition to this, GAMMs allow for the inclusion of random effects within the model. We specified cubic splines as smooth terms and set k = 4 as the knots of the spline. We also ran analyses with varying numbers of knots and created an interactive user platform to allow for the exploration of the trajectories of the metrics along the lifespan at the different numbers of knots and model parameters (Supplemental Info). Sex, scanner and intracranial volume (ICV) or total brain volume (TBV) were included as fixed effects; ages and the interaction of sex and age (sex × age) terms were subjected to the smooth kernel included to test for a sex difference in the age slope (sex × age interaction). Participant identifiers were modelled as random effects. Including the scanner as random effect did not change the results. To better appraise the sex × age interaction, estimates for age were also implemented and visualized for females and males separately. GAMMs were implemented using the mgcv R package (32). Bonferroni was used to correct for multiple comparisons in all analyses.

#### Longitudinal modeling of gyral and sulcal metrics: the association of change in gyral and sulcal features with change in cortex volume across the lifespan

In order to determine how the annual rate of change of the gyral and sulcal metrics predicted the rate of change in cortex volume across the lifespan we first determined the symmetrized annual percent change (APC) for each participant and metric *m*:

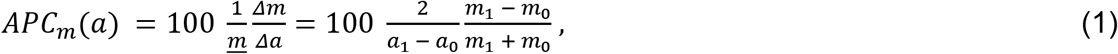

where *Δm* represents change in metric *m*, obtained at two different time points, *i.* and *i.* – 1, *i.* ∈ [0,1] and Δαthe time lapse between them in years; *a* is the participant’s age in years at a specific time point. Finally, *<u>m</u>* represents the average of the metric’s value at the two time points.

For calculation of the APC of each participant we used the value of each metric at baseline and at the nearest available follow-up measurement. We then determined the association between the APC of cortex volume and each of the APCs of the other metrics. We did this in an age-dependent manner, by linear regression of APC(age) of cortical volume on APC(age) of the gyral and sulcal metrics, locally weighted by age (Gaussian kernel, σ = 10 year) and with average TBV (across timepoints) and scanner as covariates, more detailed information about the method is given in (5). We then plotted the beta values from these regressions with their 1-standard error (SE) bands. We investigated whether there were significant sex differences in these associations by assessing overlap of the SE bands.

#### Longitudinal modeling of gyral and sulcal metrics: variability of gyral and sulcal brain metrics across the lifespan

To examine whether variability of a metric across the lifespan was constant and whether it differed between the sexes we first calculated the residuals from the GAMMs for each metric, with age, scanner, and TBV as fixed effects and participant identifiers as random effects. The quadratic value of the residuals was then used in a linear regression, as follows:

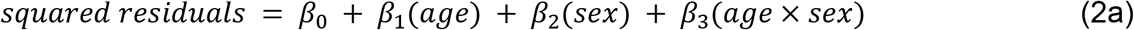

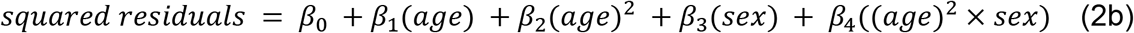

There were no quadratic effects of age and thus only results for linear age effects are presented. When there was no significant sex×age effect we ran a second version of the model not including the interaction term. If a main effect of sex or a significant effect of the sex×age interaction was found, we calculated Fisher’s variance ratio (VR) by dividing the ordinary variance of the original residuals of females by males (VR_FM_); thus, a VR_FM_ < 1 represents larger variability in males. For each outcome, a permutation test of the hypothesis that the sex specific standard deviations were equal, was performed. This was done by random permutation of the sex variable among the residuals (5000 permutations). Permuted p-values were Bonferroni corrected for multiple testing.

To characterize the differences in inter-individual variability between males and females, shift functions were estimated for each metric, using the original residuals from the GAMMs. One can characterize how two independent distributions differ by plotting the difference between the quantiles of two distributions as a function of the quantiles of one group. This technique is called a shift function and is both a graphical and inferential method (33). Wilcox proposed to use the Harrell-Davis quantile estimator to estimate the deciles of the two distributions and the computation of 95% confidence intervals of the decile differences with a bootstrap estimation of the deciles’ standard error such that the type I error rate remains around 5% across the nine confidence intervals (this means that the confidence intervals are larger than what they would be if the two distributions were compared at only one decile) (34). Quantile distribution functions were estimated for males and females separately. Let □ be a proportion (‘probability’) between 0 and 1. The quantile function specifies the metric value *X* for which a fraction *q* of the observations has metric value at or below a given *X*. The quantile function for males is given as *X_males(q)_* and for females as *X_females(q)_*. The quantile distance function is then defined as:

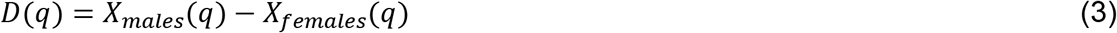

A plot of *X_males(q)_* on the x-axis vs. *D(q)* on the y-axis is then made. If *D*(*q*) is a straight-line parallel to the x-axis, this indicates a stable difference between the sexes across the distribution and thus no detectable difference in variability. A positive slope between the two extremes indicates greater male variance. A negative slope of the quantile distance function would indicate larger variability in females. The shift function was implemented using the rogme R package (35).

## Results

### Sample characteristics

No significant differences were found between males and females in age distribution, years of education, or estimated IQ (see Table 1).

### Longitudinal modeling of gyral and sulcal metrics: trajectories

We visualized age trajectories in females and males separately i) after adjusting for TBV (see Figure 1B), ii) after adjusting for ICV (see SFigure 3), and iii) not adjusting for TBV or ICV (see SFigure 4). After adjusting for TBV, all metrics displayed decrease between 16-70 years in the whole sample, except sulcal width and hull surface area, which increased with age (see Figure 1B). Trajectories of cortical thickness development were significantly different between males and females after adjusting for either ICV or TBV (see STables 1-2, Figure 1B and SFigure 3) with females having thicker cortex and a steeper decline of cortical thickness until age 40 years, approximately. There was a significant sex×age interaction for hull area after adjusting for ICV, but this interaction effect was not present after adjusting for TBV (see STables 1-2, Figure 1B and SFigure 3).

### Longitudinal modeling of gyral and sulcal metrics: the association of change in gyral and sulcal features with change in cortex volume and sulcal surface area

As can be seen in Figure 2 (rows 1-3), APC of pial surface area and cortical thickness showed the strongest associations with the APC of cortex volume, while sulcal depth and width showed the weakest associations. The APC of sulcal length showed a considerably stronger association with the APC of cortex volume as compared with depth and width. While the APC of sulcal surface area had a constant positive association with the APC of cortex volume, hull surface area displayed a profoundly different pattern across age in both sexes, i.e. a positive association with the APC of cortex volume at younger ages followed by a strong decrease in the strength of association throughout life. Across the lifespan, the APC of sulcal length was considerably more strongly associated with the APC of sulcal surface area compared with the APCs of sulcal width and depth (see Figure 2, row 4).

**Figure 2.**
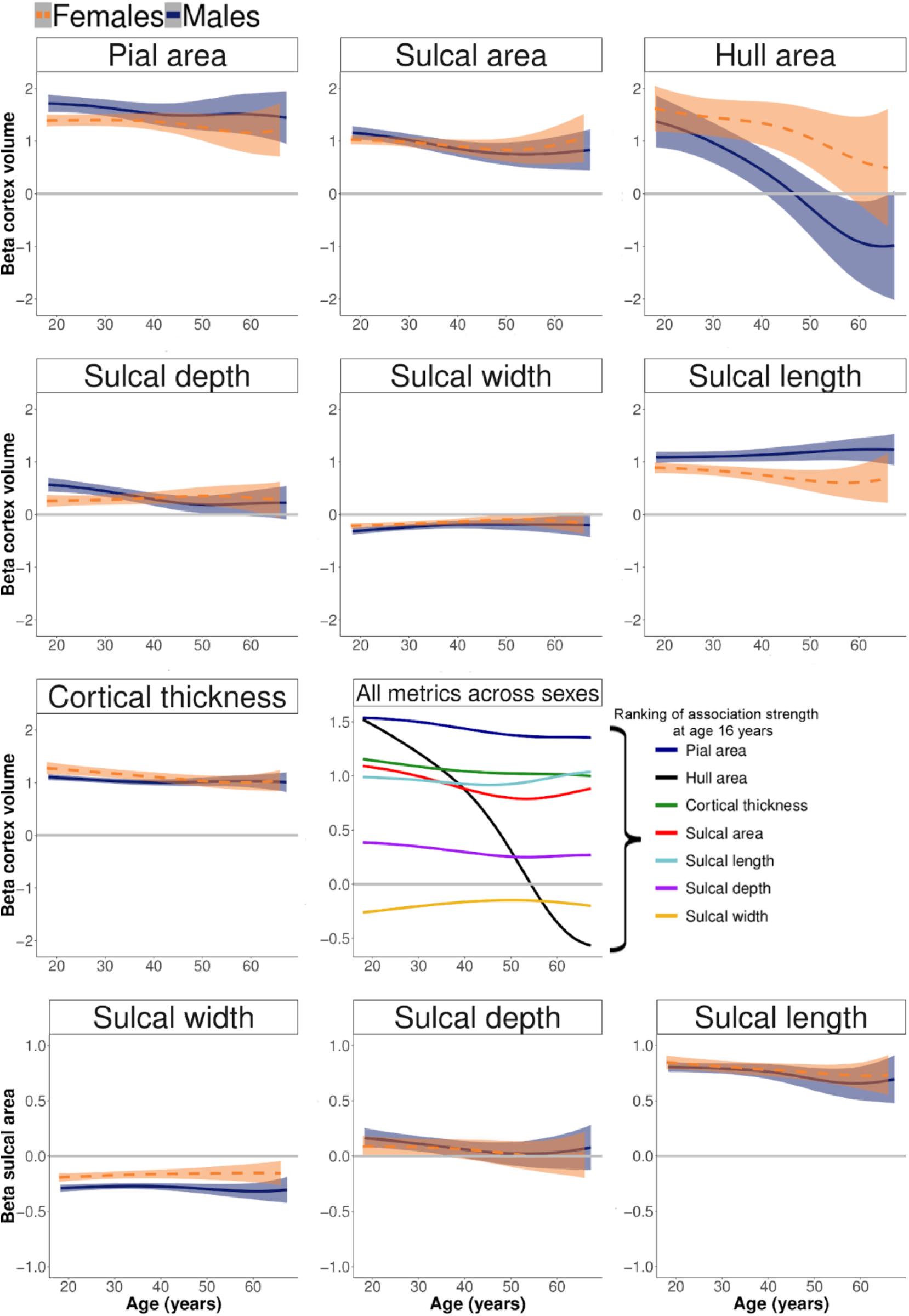
Rows 1-3: The association between the beta of symmetrized annual percent change (APC) of cortex volume and the betas of the APCs of the gyral and sulcal metrics. Row 4: The association between the beta coefficient of symmetrized APC of sulcal surface area and the betas of the APCs of sulcal depth, width and length. Beta values are from a linear regression of APC(age) of cortex volume or sulcal surface area on APC(age) of the gyral and sulcal metrics, locally weighted by age (Gaussian kernel, σ = 10 year) with average TBV (across timepoints) and scanner as covariates. The error-bands represent ± 1 SE. Solid lines represent males, dashed lines represent females. The right graph on the third row displays the association of the APC of cortex volume with the APCs of all metrics across sexes without error bands and a ranking of metrics by association strength at the youngest age (i.e. 16 years).

Sex differences in the strength of the associations (visible as a non-overlap of the 1-SE-bands) were present i) between cortex volume APC and sulcal length APC across the lifespan, and ii) between cortex volume APC and both pial area and sulcal depth APCs in young adults, with males having a higher association-strength than females for the three metrics (see Figure 2, rows 1-3). The APC of sulcal surface area was more strongly associated with the APC of sulcal width in males compared to females across the lifespan (see Figure 2, row 4).

### Longitudinal modeling of gyral and sulcal metrics: variability of gyral and sulcal brain metrics across the lifespan

After adjusting for age, scanner, TBV and the random effect of the individual, there was a main effect of sex for inter-individual variability in sulcal width (with a larger variance in females), and in cortical thickness and volume (with a larger variance in males) (see STable 4 and Figure 3). There were no sex differences in the association between inter-individual variability and age. Variance decreased with age in the whole sample for all metrics, except for cortical thickness and volume (see STable 4 and Figure 3).

**Figure 3.**
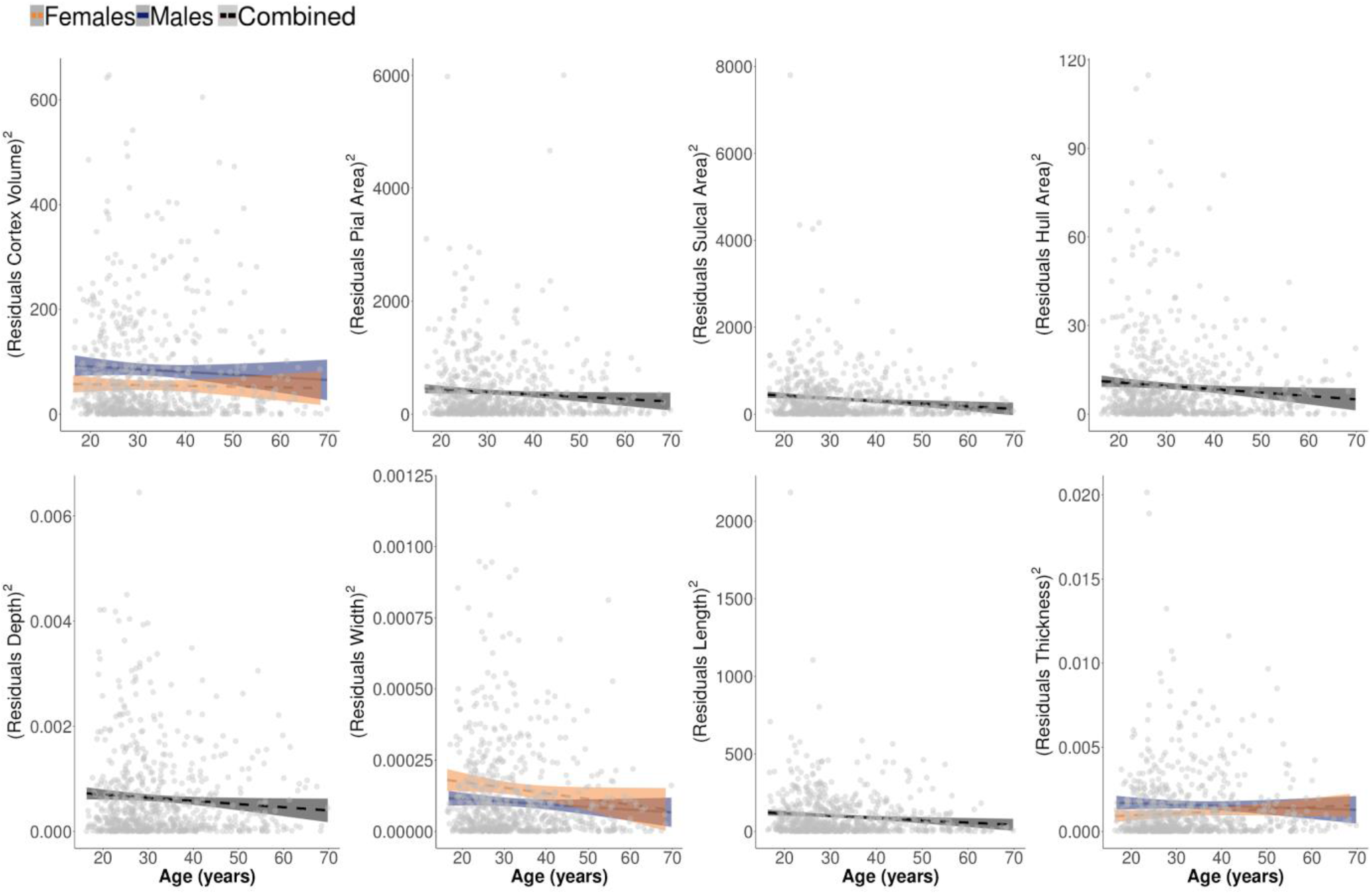
The relationship between metric inter-individual variability and age. Variability was defined as the squared residuals from the GAMMs with age, scanner, and total brain volume as fixed effects and participant identifiers as random effects. Solid lines represent males, dashed lines represent females. The age dependency of the variability was then estimated by linear regression. Across both sexes, variability decreased with age except for cortical volume and thickness (see STable 4). There was a main effect of sex for cortex volume, cortical thickness, and sulcal width and there were no significant sex×age interactions. For metrics with no main effect of sex only the regression slope of the combined sample is displayed.

The sex effects were confirmed by the VRFM with larger variability for cortex volume and cortical thickness in males (variance ratio: 0.65 and 0.70, respectively; all Bonferroni corrected permuted p-values < 0.05), and larger variability for sulcal width in females (variance ratio: 1.47, Bonferroni corrected permuted p-value < 0.05). As can be seen in Figure 4, the expression of larger variability for cortex volume in the group of males is due to a broadening of the distribution in both extremes of the distribution, while for cortical thickness this was more prominent in the lower (left extreme) deciles. For sulcal width, females had larger variability in the lower (left extreme) deciles.

**Figure 4.**
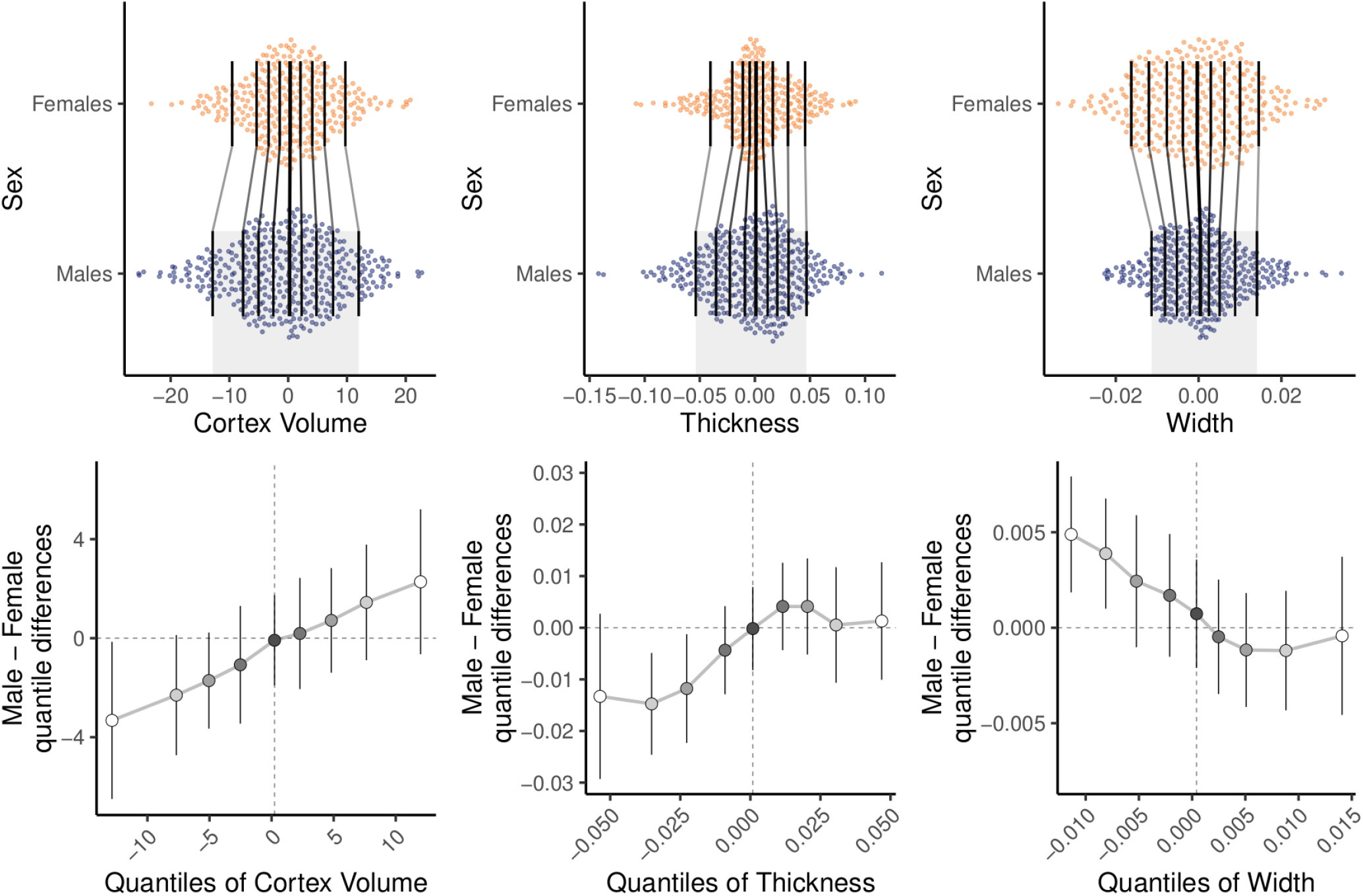
Marginal distributions for cortex volume, cortical thickness and sulcal width for males and females after adjusting for age, scanner, total brain volume and the random effect of the individual. Gray lines show the amount of shift between the two distributions (35). A sloped line indicates a difference in the distributions between the groups. In the bottom row, the magnitude of the sex difference is plotted as a function of the distribution among males. A positively sloped grey line indicates larger variability in males. Error bars represent bootstrapped 95% CIs (5000 bootstraps).

## Discussion

In this longitudinal study we investigated how global sulcal and gyral metrics of cortex morphology change with age in individuals between 16 and 70 years of age and explored whether there are sex differences in the lifespan trajectories of these metrics. We also examined the contribution of change in gyral and sulcal metrics to cortical volume loss across the lifespan and the effect of sex and age on the strength of the association between these metrics and cortex size. Finally, we assessed whether sex influenced the age-dependency of the inter-individual variability of these metrics.

### Developmental trajectories of gyral and sulcal morphology

After adjusting for total brain volume, we found that sulcal surface area decreases from adolescence through late adulthood, while hull surface area does not, and, in fact, even shows a slight increase with age. The finding of sulcal surface area loss and gyral surface area conservation across the lifespan extends a previous finding in an independent sample of smaller loss of hull surface area compared to overall cortical surface area loss during late adolescence (10). We found a significant effect of age on the three subcomponents of sulcal surface area: sulcal depth, width and length, with no significant sex differences in trajectories for these metrics. In the whole sample, shallowing of sulci takes place at a constant pace, widening of sulci levels off between 30-40 years, but resumes thereafter at a constant rate, and shortening of sulci is most severe before 40 years. To our knowledge, the current study is the first to show the dynamic changes in sulcal length across the lifespan in both sexes separately. It also confirms prior cross-sectional studies reporting age-related decreases in sulcal depth and increases in sulcal width in the human brain (11,36–38). Sulcal shortening may be reflective of a decrease in sulcal crookedness rather than disappearance of sulci (39,40). Sulcal shallowing and widening over time may be the result of shrinkage in the adjacent gyri but may also result from more distal changes in grey and white matter and, combined with sulcal shortening, may result in a loss of sulcal surface area (37,41,42).

### Effect of sex on metrics and developmental trajectories of gyral and sulcal morphology

Across the lifespan, we found sex differences in some gyral and sulcal metrics. The female group had thicker cortex and deeper sulci than the male group, which is consistent with previous reports in younger and older populations (43,44). We also found trajectories of cortical thickness development that differed between the sexes. At the youngest age, the female group shows a thicker cortex and this sex difference appears to decrease until it disappears at age 40 years approximately. Although widespread thicker cortex in young females has been previously reported, recent longitudinal studies in children and adolescents have yielded mixed results, with some reports only showing localized increases of cortical thickness in females and no sex differences in cortical thickness trajectories (45,46). The cause for thicker cortex in younger females is unclear but it may be the result of puberty and the related changes in sex hormones, such as estrogens, which may have a neuro-protective effect on brain structure (47,48). Interestingly, a potential neuroprotective effect of estrogens has been considered to potentially underlie the second incidence peak of schizophrenia that may be found in females after the age of 40, approximately the same age where we find an attenuation of sex differences in cortical thickness measurements (49,50). Reduced cortical thickness and progressive thinning has been consistently found in schizophrenia and they seem to be associated with worsening of clinical symptoms and prognosis (20,51). It remains to be elucidated whether biological mechanisms underpinning differing trajectories of cortical thickness development in healthy females and males may also underlie some of the sex differences in the epidemiology and clinical presentation of psychotic and other mental disorders (52,53).

### The association of change in gyral and sulcal features with change in cortex volume across the lifespan

We found a steep decline in the strength of association between changes in hull area and cortex volume across the age range studied in both sexes. Thus, in our sample loss of cortex volume co-occurs with loss of hull surface area during late adolescence and early adulthood, but the strength of this association decreases with age and, in males, the direction of the association tends to invert in late adulthood. The declining contribution of hull surface area loss to cortex volume loss suggests that overall loss of pial surface area across the lifespan is primarily driven by loss of sulcal surface area across the study’s age range. Sulcal surface area makes up the majority of the pial surface. While sulcal surface area decreases as it becomes wider, shallower and shorter, the surface surrounding the gyral crown may be preserved and possibly stretched, thus eventually leading to the inverse relationship with cortex volume in males at later stages.

Across the sexes and the age range studied, the strongest contributor to the decrease in sulcal surface area is sulcal shortening, as change in sulcal length had the strongest relationship with change in sulcal surface area across the lifespan. This finding extends a reported strong cross-sectional association between sulcal length and sulcal surface in young people aged 12-13 years (16). Apart from sulcal crookedness, sulcal length is also dependent on the number of sulci. However, given that disappearance of sulci during adolescence and adulthood is highly unlikely, our results rather manifest that cortical volume loss over time may be more due to a decrease in sulcal tortuosity over time compared to sulcal widening or shallowing and that this effect is stronger in males than in females. Sexual dimorphism in the patterns of association between gyral and sulcal metrics and cortex size, with greater effects found in males at older ages, may reflect a process of accelerated brain aging in late adulthood in healthy males relative to females, as suggested by some cognitive and brain metabolic data (54,55).

### Inter-individual variability of morphological metrics

We found a significant effect of sex on inter-individual variability across the lifespan in several brain metrics. Males showed greater variability in cortex volume and cortical thickness, thus replicating cross-sectional studies in lifespan and older samples. In the current study males show a wider distribution, towards both tails, for cortex volume when compared to females. For cortical thickness in males and for sulcal width in females, we found greater variability for the lower values of the distribution. It has been hypothesized that increased male variability in cortical volume and thickness is related to the observation that in species with sex chromosomes and sex chromosome dosage compensation (i.e. the compensatory mechanisms that balance gene expression between autosomes and sex chromosomes in the heterogametic sex), heterogametic individuals tend to exhibit greater phenotypic variance (the “sex-chromosome hypothesis”) (56). Increased variability of cortical thickness and volume in sex-biased mental disorders such as schizophrenia and autism spectrum disorders (with increased prevalence in males) indirectly support this claim (57,58). In these disorders greater inter-individual variability of volume and cortical thickness increases the probability of extremely deviating individuals towards the ends of the distribution. Characterizing these deviating individuals is clinically important as they may drive group average findings and constitute a cluster with a separate phenotypic presentation (59,60).

Our findings of increased relative variability of sulcal width among females confirm the findings of a study that assessed sulcal width of ten primary sulci and found that while the male group showed greater sulcal width overall, the standard deviations for all sulci were higher in the female group (11). Given the lower heritability of global sulcal width compared to cortical thickness and volume, behavioral and environmental factors may have a larger influence on variability of sulcal width (27,61). Nevertheless, whether and how genetic and non-genetic factors (including the effects of gender in the context of gendered societies) contribute to sex differences in variability of brain structure is something that future studies including morphological, genetic and environmental information should address.

To the best of our knowledge, this is the first study looking at sex differences in age trajectories of inter-individual variability of sulcal morphological measurements. We found no sex differences in the age-dependency of morphological variation. Across sexes, relative variance of these metrics decreased with age, except for cortical thickness and volume, which is consistent with a recent cross-sectional lifespan study reporting either larger morphological variation in younger than older healthy individuals or no age-dependency of variability (62).

The results of this study should be interpreted with caution due to the following limitations. First, there are fewer participants at both extremes of the age distribution, which is reflected in larger confidence intervals. Second, since the current sample consisted of control participants in a longitudinal clinical study, they were extensively screened for clinical conditions. One may speculate whether this group is representative of the broader population across the studied age range. However, we replicate a number of findings from other studies which lead us to believe that our population fits well with the other populations described in the literature on longitudinal changes in brain morphology. Third, we focus on global measurements instead of assessments at the level of regions or individual sulci. However, by focusing on global measurements, this study provides an empirical and methodological basis for future studies assessing local sulcal morphology. Fourth, this study cannot completely separate the effects of sex from gender, which also influences exposure to environmental and social factors across the lifespan and could underlie, at least partially, some of the reported differences in brain structure between sexes.

This study offers a novel framework for prospective longitudinal studies of gyral and sulcal morphology in health and disease. Our findings support a differential effect of age and sex on longitudinal trajectories of several metrics of gyral and sulcal morphology across the lifespan and suggest that change in sulcal surface area, notably through sulcal shortening, may be a main contributor to change in cortex volume associated with aging across sexes. The study of the genetic, environmental and social determinants underlying sex differences in cortical development and aging may provide invaluable information about typical brain development and the pathways into psychiatric and neurological conditions.

## Acknowledgements

This work was supported by the Spanish Ministry of Science and Innovation, Instituto de Salud Carlos III (PI17/01249, PI17/00997, PI19/01024), co-financed by ERDF funds from the European Commission, “A way of making Europe”, CIBERSAM, Madrid Regional Government (B2017/BMD-3740 AGES-CM-2), European Union Structural Funds, European Union Seventh Framework Program under grant agreements FP7-HEALTH-2009-2.2.1-2-241909 (Project EU-GEI), FP7-HEALTH-2009-2.2.1-3-242114 (Project OPTiMISE), FP7-HEALTH-2013-2.2.1-2-603196 (Project PSYSCAN) and FP7-HEALTH-2013-2.2.1-2-602478 (Project METSY); and European Union H2020 Program under the Innovative Medicines Initiative 2 Joint Undertaking (grant agreement No 115916, Project PRISM, and grant agreement No 777394, Project AIMS-2-TRIALS), Fundación Familia Alonso, Fundación Alicia Koplowitz, and Fundación Mutua Madrileña. Dr. Díaz-Caneja holds a Juan Rodés grant from Instituto de Salud Carlos III (JR19/00024).

The authors thank Zimbo Boudewijns, Joyce van Baaren and Diego Muñoz Beltrán for technical assistance.

