## Supplement for "Sex differences in lifespan trajectories and variability of human sulcal and gyral morphology"

*Image quality assessment*

After the completion of the preprocessing pipeline for all T1-weighted MRI scans in FreeSurfer, we rigorously assessed the quality of the data using a combination of visual inspection and the examination of several quantitative quality assessment (QA) metrics. First, we calculated five QA measurements based on those proposed by the Preprocessed Connectome Project (http://preprocessed-connectomes-project.org/quality-assessment-protocol/#spatial-anatomical) to identify images that were unusable: signal-to-noise ratio (SNR), contrast to noise ratio (CNR), foreground to background energy ratio (FBER), percent artefact voxels (Artifacts) and entropy focus criterion (Entropy). Following ENIGMA criteria (http://enigma.ini.usc.edu/protocols/imaging-protocols/) we defined the threshold for outliers as [mean-(2.698*SD)] for SNR, CNR and FBER metrics; and as [mean+(2.698*SD)] for Artifacts and Entropy. Next, we calculated for each scan mean CT, total SA, total WM volume, total GM volume, subcortical GM volume and ICV for the whole brain. Then, we summed for each image the amount of outliers over all measures, calculated the mean+(2.698*SD) for the amount of outliers over the whole sample, designated those above the threshold as outliers, and subsequently removed them from further analysis.

*QA-based exclusions*

After the evaluation of the computed quantitative measures of image quality, 138 scans were considered of insufficient quality and were excluded from our analysis (see SFigure 1A). These excluded scans were validated by manually assessing image quality after preprocessing to ensure that all preprocessing steps worked and to avoid the dissemination of error along the analysis. The parameters that were most useful to objectively detect artefacts visually were agreed on between researchers, e.g. incorrect sulcal labelling or insufficient quality of sulcal segmentation resulting in gross anatomical abnormalities. These visual checks were performed in BrainVisa. Those scans with poor initial sulcal labelling were manually edited, which often resulted in better sulcal labelling when manually inspected (see SFigure 1B) for examples of "good", "medium" and "bad" quality sulcal labelling after first preprocessing of the images). Twelve additional scans were excluded after this manual inspection.

SFigure 1.

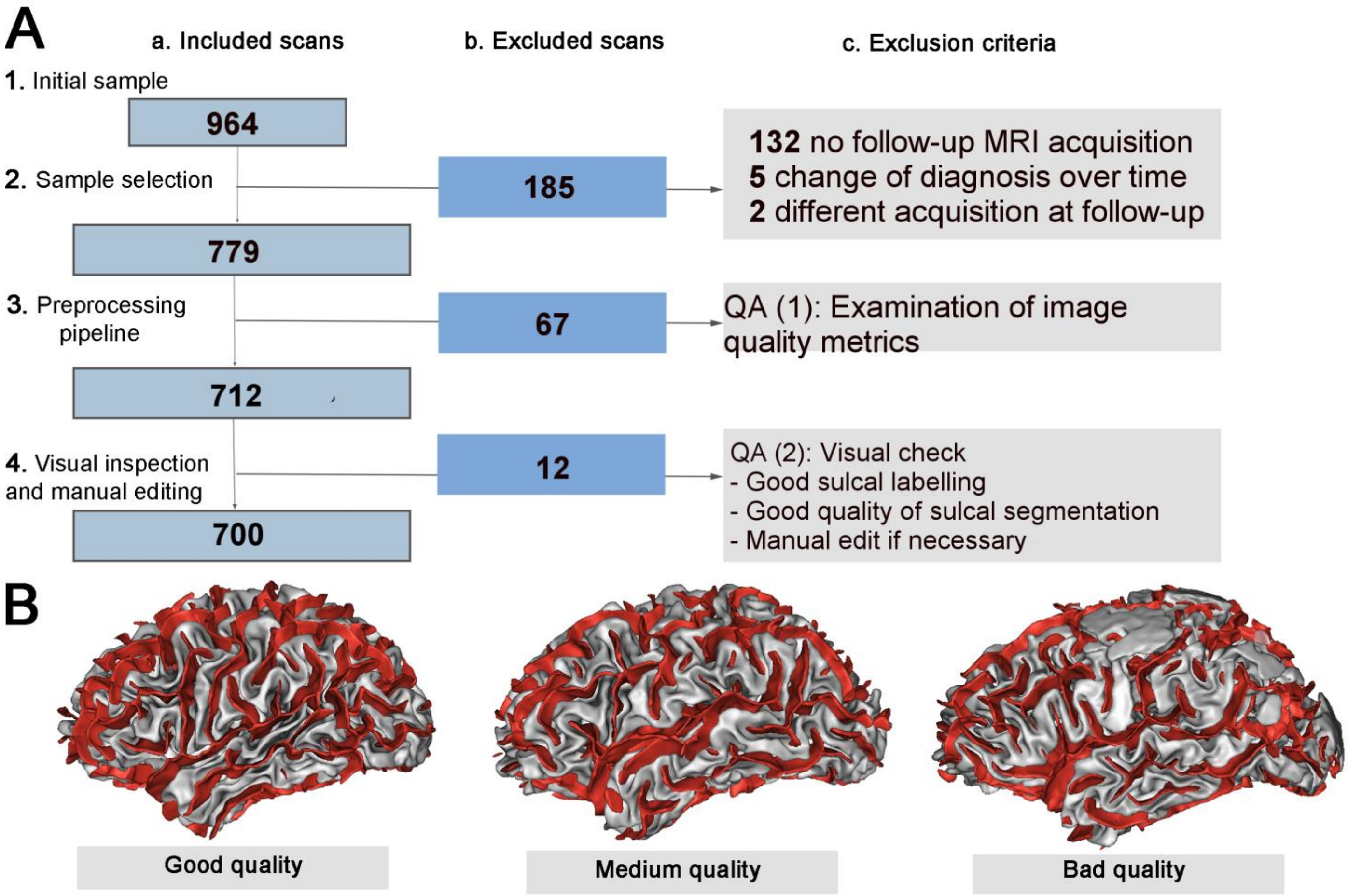

SFigure 2. Explained variance for age, sex and scanner at baseline (**A**) and first follow-up (**B**) for each brain structural metric. Each participant was always scanned on the same scanner. For **A** and **B**: The left column displays the percentage of variance explained by age, sex and scanner for each metric. The right column shows that, across metrics, sex and age were the dominant sources of variance explaining on average about 20% and 14% of the total variance at baseline and follow-up, respectively, while scanner explained on average less than 1% of the variance at baseline and follow-up.

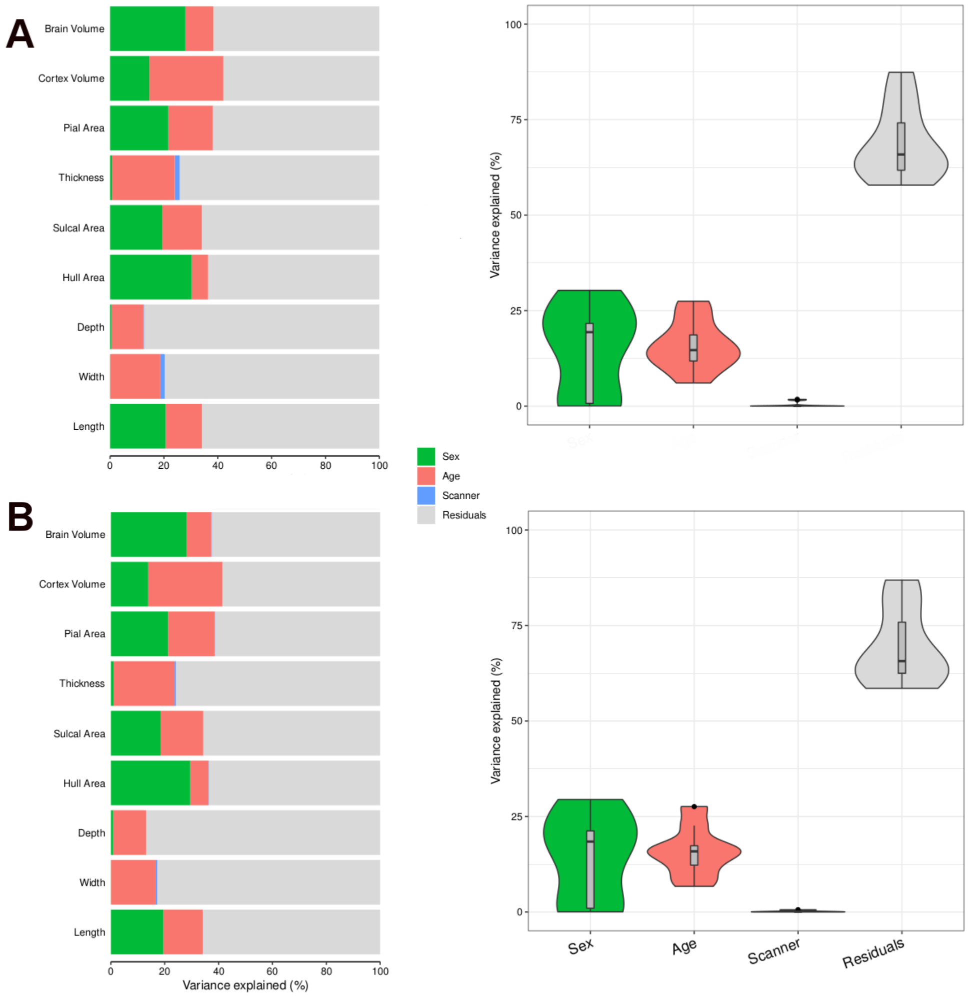

SFigure 3. Generalized additive mixed model (GAMMs) fits for males (blue) and females (orange). Fits are shown with confidence intervals. Intracranial volume and scanner are included in the models as covariates.

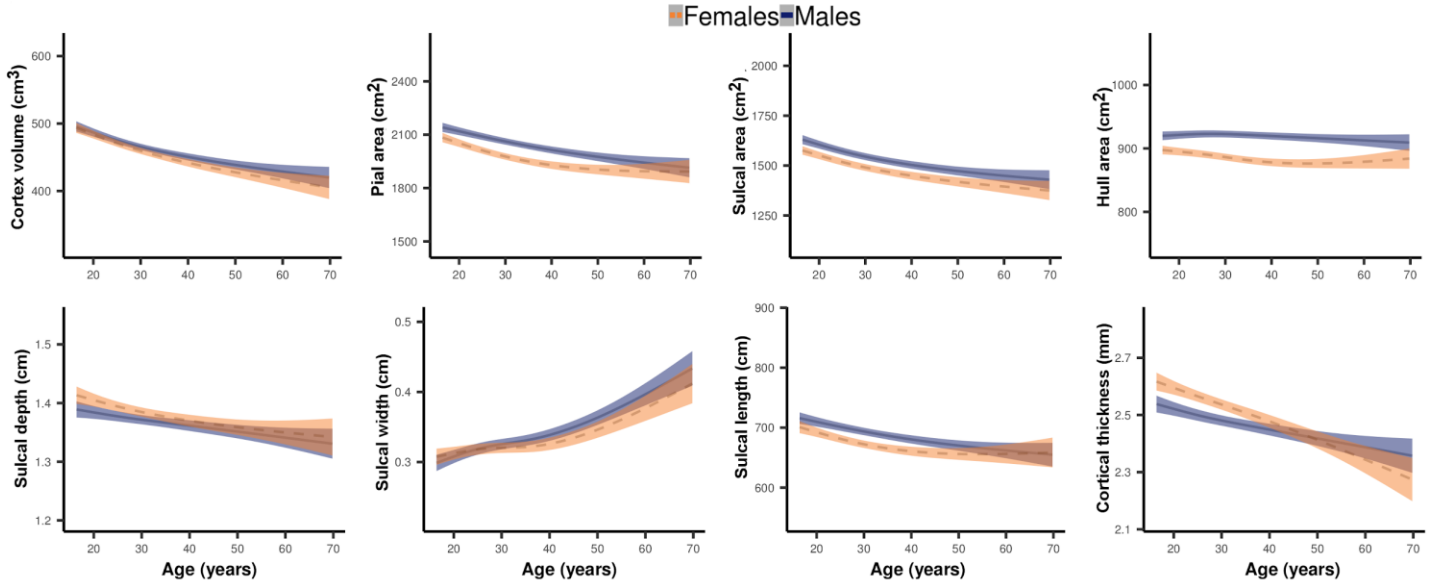

SFigure 4. Generalized additive mixed model (GAMMs) fits for males (blue) and females (orange). Fits are shown with confidence intervals over the raw data points for all participants in the corresponding color, with data at each of the time points connected for each participant. Scanner was included in the models as a covariate.

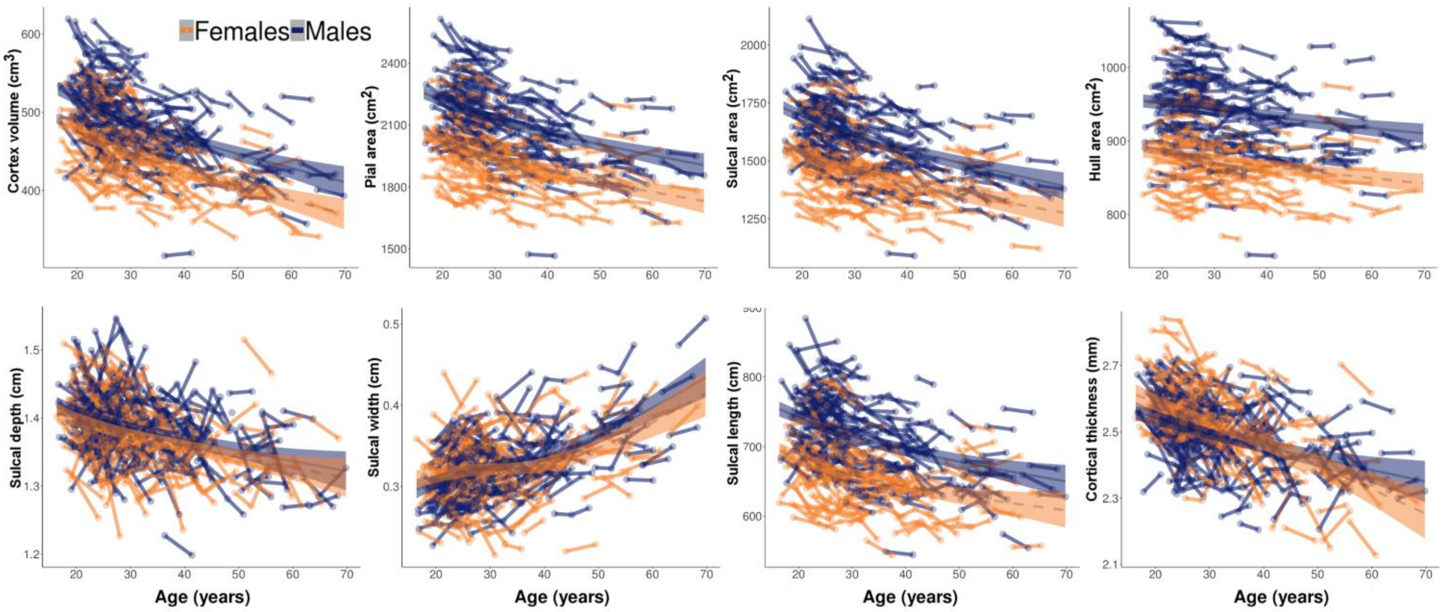

STable 1. Generalized additive mixed model (GAMMs) estimates for sex, age, and sex*age for cortex volume, pial surface area, sulcal surface area, hull surface area, sulcal depth, sulcal width, sulcal length, and cortical thickness. We included scanner and total brain volume as covariates and participant identifiers as a random effect in the models. Smooth function (edf), degrees of freedom (Ref.df), F-statistic, and significance are given (p-values in bold are significant after Bonferroni correction).

| **Brain metrics** | **GAMM estimates** | | | |
|---|---|---|---|---|
| Cortex Volume | | | | |
| Intercept | Estimate | SE | t | p |
| sex | -6.46 | 2.61 | -2.48 | **0.01** |
| Slope | edf | Ref.df | F | p |
| s(age) | 2.59 | 3 | 1369.09 | **<0.001** |
| s(age):males | 1.85 | 3 | 51.69 | 0.03 |
| Pial Area | | | | |
| Intercept | Estimate | SE | t | p |
| sex | 22.32 | 10.25 | 2.18 | 0.03 |
| Slope | edf | Ref.df | F | p |
| s(age) | 2.75 | 3 | 4137.61 | **<0.001** |
| s(age):males | 0.11 | 3 | 0.82 | 0.27 |
| Sulcal Area | | | | |
| Intercept | Estimate | SE | t | p |
| sex | 4.36 | 10.68 | 0.41 | 0.68 |
| Slope | edf | Ref.df | F | p |
| s(age) | 2.75 | 3 | 3491.85 | **<0.001** |
| s(age):males | 0.57 | 3 | 37.11 | 0.15 |

| Hull Area | | | | |
|---|---|---|---|---|
| Intercept | Estimate | SE | t | p |
| sex | 11.94 | 2.29 | 5.21 | **<0.001** |
| Slope | edf | Ref.df | F | p |
| s(age) | 2.77 | 3 | 1804.48 | **0.002** |
| s(age):males | 1.42 | 3 | 737.76 | 0.03 |
| Depth | | | | |
| Intercept | Estimate | SE | t | p |
| sex | -0.02 | 0.01 | -3.27 | **0.001** |
| Slope | edf | Ref.df | F | p |
| s(age) | 1.83 | 3 | 101.99 | **<0.001** |
| s(age):males | 0 | 3 | 0 | 0.46 |
| Width | | | | |
| Intercept | Estimate | SE | t | p |
| sex | 0.03 | 0.01 | 5.63 | **<0.001** |
| Slope | edf | Ref.df | F | p |
| s(age) | 2.73 | 3 | 806.84 | **<0.001** |
| s(age):males | 1.62 | 3 | 154.17 | 0.05 |
| Length | | | | |

| Intercept | Estimate | SE | t | p |
|---|---|---|---|---|
| sex | 6.9 | 4.01 | 1.72 | 0.09 |
| Slope | edf | Ref.df | F | p |
| s(age) | 2.47 | 3 | 599.56 | **<0.001** |
| s(age):males | 0.57 | 3 | 14.97 | 0.13 |
| Thickness | | | | |
| Intercept | Estimate | SE | t | p |
| sex | -0.06 | 0.01 | -4.37 | **<0.001** |
| Slope | edf | Ref.df | F | p |
| s(age) | 2.26 | 3 | 1335.4 | **<0.001** |
| s(age):males | 2.04 | 3 | 215.5 | **<0.001** |

STable 2. Generalized additive mixed model (GAMMs) estimates for sex, age, and sex x age for cortex volume, pial surface area, sulcal surface area, hull surface area, sulcal depth, sulcal width, sulcal length, and cortical thickness. We included scanner and intracranial volume as covariates and participant identifiers as a random effect in the models. Smooth function (edf), degrees of freedom (Ref.df), F-statistic, and significance are given (p-values in bold are significant after Bonferroni correction).

| **Brain metrics** | **GAMM estimates** | | | |
|---|---|---|---|---|
| Cortex Volume | | | | |
| Intercept | Estimate | SE | t | p |
| sex | 6.63 | 3.1 | 2.14 | 0.030 |
| Slope | edf | Ref.df | F | p |
| s(age) | 2.59 | 3 | 1951.07 | **<0.001** |
| s(age):males | 0.69 | 3 | 16.31 | 0.100 |
| Pial Area | | | | |
| Intercept | Estimate | SE | t | p |
| sex | 78.93 | 12.3 | -6.34 | **<0.001** |
| Slope | edf | Ref.df | F | p |
| s(age) | 2.61 | 3 | 12045.54 | **<0.001** |
| s(age):males | 0 | 3 | 0 | 0.570 |
| Sulcal Area | | | | |
| Intercept | Estimate | SE | t | p |
| sex | 54.12 | 11.89 | 4.55 | **<0.001** |
| Slope | edf | Ref.df | F | p |
| s(age) | 2.59 | 3 | 9281.05 | **<0.001** |
| s(age):males | 0 | 3 | 0 | 0.550 |

| | | | | |
|---|---|---|---|---|
| Hull Area | | | | |
| Intercept | Estimate | SE | t | p |
| sex | 33.33 | 3.48 | 9.58 | **<0.001** |
| Slope | edf | Ref.df | F | p |
| s(age) | 2.57 | 3 | 5068.29 | **<0.001** |
| s(age):males | 2.48 | 3 | 1631.16 | 0.010 |
| Depth | | | | |
| Intercept | Estimate | SE | t | p |
| sex | -0.013 | 0.01 | -2.5 | 0.010 |
| Slope | edf | Ref.df | F | p |
| s(age) | 1.89 | 3 | 122.24 | **<0.001** |
| s(age):males | 0 | 3 | 0 | 0.310 |
| Width | | | | |
| Intercept | Estimate | SE | t | p |
| sex | 0.01 | 0.01 | 1.2 | 0.230 |
| Slope | edf | Ref.df | F | p |
| s(age) | 2.7 | 3 | 1071.14 | **<0.001** |
| s(age):males | 1.41 | 3 | 186.81 | 0.010 |
| Length | | | | |

| Intercept | Estimate | SE | t | p |
|---|---|---|---|---|
| sex | 18.55 | 4.16 | 4.46 | **<0.001** |
| Slope | edf | Ref.df | F | p |
| s(age) | 2.35 | 3 | 1735.69 | **<0.001** |
| s(age):males | 0.01 | 3 | 0 | 0.380 |
| Thickness | | | | |
| Intercept | Estimate | SE | t | p |
| sex | -0.05 | 0.01 | -3.7 | **<0.001** |
| Slope | edf | Ref.df | F | p |
| s(age) | 2.3 | 3 | 1389.04 | **<0.001** |
| s(age):males | 1.8 | 3 | 186.29 | **<0.001** |

STable 3. Generalized additive mixed model (GAMMs) estimates for sex, age, and sex*age for cortex volume, pial surface area, sulcal surface area, hull surface area, sulcal depth, sulcal width, sulcal length, and cortical thickness. Models are not adjusted for ICV or TB; we included scanner as a covariate and participant identifiers as a random effect in the models. Smooth function (edf), degrees of freedom (Ref.df), F-statistic, and significance are given (p-values in bold are significant after Bonferroni correction).

| **Brain metrics** | **GAMM estimates** | | | |
|---|---|---|---|---|
| Cortex Volume | | | | |
| Intercept | Estimate | SE | t | p |
| sex | 40.97 | 4.61 | 8.89 | **<0.001** |
| Slope | edf | Ref.df | F | p |
| s(age) | 2.72 | 3 | 21392.01 | **<0.001** |
| s(age):males | 0 | 3 | 0 | 0.45 |
| Pial Area | | | | |
| Intercept | Estimate | SE | t | p |
| sex | 188.37 | 17.97 | 10.48 | **<0.001** |
| Slope | edf | Ref.df | F | p |
| s(age) | 2.76 | 3 | 54899.89 | **<0.001** |
| s(age):males | 0.63 | 3 | 254.56 | 0.120 |
| Sulcal Area | | | | |
| Intercept | Estimate | SE | t | p |
| sex | 151.25 | 16.16 | 9.36 | **<0.001** |
| Slope | edf | Ref.df | F | p |
| s(age) | 2.71 | 3 | 25523.8 | **<0.001** |
| s(age):males | 1.05 | 3 | 763.5 | 0.030 |

| | | | | |
|---|---|---|---|---|
| Hull Area | | | | |
| Intercept | Estimate | SE | t | p |
| sex | 67.14 | 5.61 | 11.97 | **<0.001** |
| Slope | edf | Ref.df | F | p |
| s(age) | 2.35 | 3 | 58901 | **<0.001** |
| s(age):males | 0 | 3 | 0 | 0.790 |
| Depth | | | | |
| Intercept | Estimate | SE | t | p |
| sex | 0.01 | 0.01 | 1.92 | 0.060 |
| Slope | edf | Ref.df | F | p |
| s(age) | 2.1 | 3 | 394.07 | **<0.001** |
| s(age):males | 0 | 3 | 0 | 0.670 |
| Width | | | | |
| Intercept | Estimate | SE | t | p |
| sex | 0 | 0 | 0.85 | 0.400 |
| Slope | edf | Ref.df | F | p |
| s(age) | 2.7 | 3 | 1130.35 | **<0.001** |
| s(age):males | 1.48 | 3 | 212.71 | 0.008 |
| Length | | | | |

| Intercept | Estimate | SE | t | p |
|---|---|---|---|---|
| sex | 54.86 | 5.58 | 9.83 | **<0.001** |
| Slope | edf | Ref.df | F | p |
| s(age) | 2.48 | 3 | 6527.03 | **<0.001** |
| s(age):males | 0.91 | 3 | 255.88 | 0.040 |
| Thickness | | | | |
| Intercept | Estimate | SE | t | p |
| sex | -0.02 | 0.01 | -2.17 | 0.030 |
| Slope | edf | Ref.df | F | p |
| s(age) | 2.31 | 3 | 1573.11 | **<0.001** |
| s(age):males | 1.83 | 3 | 152.95 | **0.002** |

STable 4. Results from linear regressions using squared residuals from generalized additive mixed models (p-values in bold are significant after Bonferroni correction).

| **Brain metrics** | **Intercept** | **SE** | **p** | **Age** | **SE** | **p** | **Sex** | **SE** | **p** |
|---|---|---|---|---|---|---|---|---|---|
| Cortex Volume | 95.92 | 12.40 | **<0.001** | -0.36 | 0.35 | 0.300 | -29.31 | 7.55 | **<0.001** |
| Pial Area | 522.53 | 74.76 | **<0.001** | -4.24 | 2.08 | 0.040 | -10.45 | 45.51 | 0.820 |
| Sulcal Area | 546.63 | 69.27 | **<0.001** | -6.08 | 1.93 | **0.002** | -5.00 | 42.16 | 0.910 |
| Hull Area | 12.49 | 1.78 | **<0.001** | -0.11 | 0.05 | 0.020 | 1.11 | 1.08 | 0.300 |
| Depth | <0.001 | 0.00 | **<0.001** | <0.001 | <0.001 | 0.040 | <0.001 | <0.001 | 0.970 |
| Width | <0.001 | 0.00 | **<0.001** | <0.001 | <0.001 | 0.020 | <0.001 | <0.001 | **<0.001** |
| Length | 148.83 | 18.33 | **<0.001** | -1.48 | 0.51 | **0.004** | -8.15 | 11.16 | 0.470 |
| Thickness | 0.00 | 0.00 | **<0.001** | <0.001 | <0.001 | 0.920 | <0.001 | <0.001 | **0.003** |
